# Tempora: cell trajectory inference using time-series single-cell RNA sequencing data

**DOI:** 10.1101/846907

**Authors:** Thinh N. Tran, Gary D. Bader

## Abstract

Single-cell RNA sequencing (scRNAseq) can map cell types, states and transitions during dynamic biological processes such as development and regeneration. Many trajectory inference methods have been developed to order cells by their progression through a dynamic process. However, when time series data is available, these methods do not consider the available time information when ordering cells and are instead designed to work only on a single scRNAseq data snapshot. We present Tempora, a novel cell trajectory inference method that orders cells using time information from time-series scRNAseq data. In performance comparison tests, Tempora accurately inferred developmental lineages in human skeletal myoblast differentiation and murine cerebral cortex development, beating state of the art methods. Tempora uses biological pathway information to help identify cell type relationships and can identify important time-dependent pathways to help interpret the inferred trajectory. Our results demonstrate the utility of time information to supervise trajectory inference for scRNA-seq based analysis.

## BACKGROUND

Dynamic tissue-level processes, such as development and regeneration, are critical for multicellular organisms. Single-cell RNA sequencing (scRNAseq) now enables us to map the range of cell types and states in these processes at cellular resolution^1^. A single scRNA-seq snapshot can be used to infer lineage relationships between cell types and states^2^. Snapshot scRNAseq studies have been used to investigate multiple aspects of development, including the early embryo, blood, differentareas of the brain and more^3^. Even though snapshot scRNAseq can provide novel insights into development, it has recognized limits^4^, including that cell populations that appear earlier or later than the sampling time cannot be studied. Time-series scRNAseq can address some of these limits and has been increasingly applied to study tissue development, including cerebral cortex^5^, kidney^6^, heart^7^ and more.

When using scRNAseq to study dynamic processes, whether through snapshot or time-series experiments, it is of interest to order cells at different stages along an axis that represents how far along they are on the process under study based on their transcriptional signatures. The ordering problem, commonly termed pseudotime ordering if it is inferred from data without a known temporal ordering, consists of two main parts: the identification of a trajectory representing the paths that cells follow, and the determination of pseudotime values for individual cells along this trajectory. This inferred trajectory enables us to study the sequential changes of gene expression during a process, as well as identify branches and instrumental genes at the branching points. More than 70 computational methods to order cells along pseudotemporal axes, known as trajectory inference methods, have been published, which employ different strategies to infer lineage and order cells^8^. Most trajectory inference methods are developed based on the basic premise that cells closer in developmental time have more similar gene expression signatures, thus a likely trajectory is a path that maximizes cell-to-cell similarity. Common strategies for trajectory construction include fitting a minimum spanning tree (MST), which connects all data points in a path that minimizes distance between points, or nonlinear dimensionality reduction that identifies a low-dimensional manifold that cells lie on. Monocle, the pioneering trajectory inference method, constructs a MST connecting all cells in a reduced dimension space, then determines the longest path through this tree as the backbone and orders cells along this path^9^. Some methods, including TSCAN and Slingshot, build the MST on cluster centers, representing cell types and states, instead of individual cells, and project cells on the MST to determine their pseudotime values^10,11^. Other methods, such as PAGA and StemID, use graph theory methods, such as graph partition, to construct trajectories^12,13^. The majority of available trajectory inference methods have been evaluated and integrated in Dyno, a platform that enables users to conveniently apply selected methods to their data^8^.

While many scRNAseq trajectory inference methods exist, none have been designed to consider time-series information. Time-series analysis of gene expression data has been studied, but mostly before the advent of high-throughput single cell transcriptomics methods and focused primarily on bulk RNAseq analysis^14^. Since cell types and states identified primarily in earlier time points must be earlier in the trajectory than those identified primarily in later time points, we hypothesize that explicit use of temporal ordering information available in time-series scRNAseq data can be used to improve trajectory inference. To address this, we introduce Tempora, a novel method to infer cell lineage maps from time-series scRNAseq data. Tempora aligns cell types and states across time points using available batch and data set alignment methods, as well as biological pathway information, then infers trajectory relationships between these cell types using the available temporal ordering information. Evaluating Tempora on two different time-series scRNAseq data sets using gold standards showed that our method outperforms established trajectory inference methods.

## RESULT

### Method overview

The Tempora method infers cell type-based trajectories from time-series scRNAseq data. Tempora focuses on identifying how cell types are related across the entire time-series data set, based on the established assumption that cells with similar gene expression profiles are closer in the cell lineage. After identifying cell type transcriptome similarity relationships, Tempora orders these links based on the time-series data. Cells identified primarily in earlier time points are ordered earlier in the trajectory than those identified primarily in later time points. To build a more robust trajectory, less influenced by small outlier populations, Tempora first clusters cells with similar transcriptional signatures and infers a trajectory that connects cell clusters rather than individual cells. These clusters represent putative cell types, such as progenitors, immune cells, cardiomyocytes, or stable cell states (e.g. cycling, proliferating or metabolizing^2^). Second, to improve robustness of the trajectory relationship identification step, clusters are compared to each other based on their biological pathway enrichment profiles instead of individual gene expression profiles. This also helps improve biological interpretability of the trajectory result, as trajectory related pathway expression patterns can be automatically identified.

Tempora takes as input a preprocessed gene expression matrix from a time-series scRNAseq experiment and cluster labels for all cells. Tempora then calculates the average gene expression profiles, or centroids, of all clusters before transforming the data from gene expression space to pathway enrichment space using single-sample gene set variation analysis (GSVA)^15^ (Figure 1). To focus on high variance and non-redundant pathway information, Tempora applies PCA on the pathway enrichment analysis result and selects important PCs using a scree plot. Pathways with high loadings on those PCs are used to construct the lineage in the next step.

**Figure 1.**
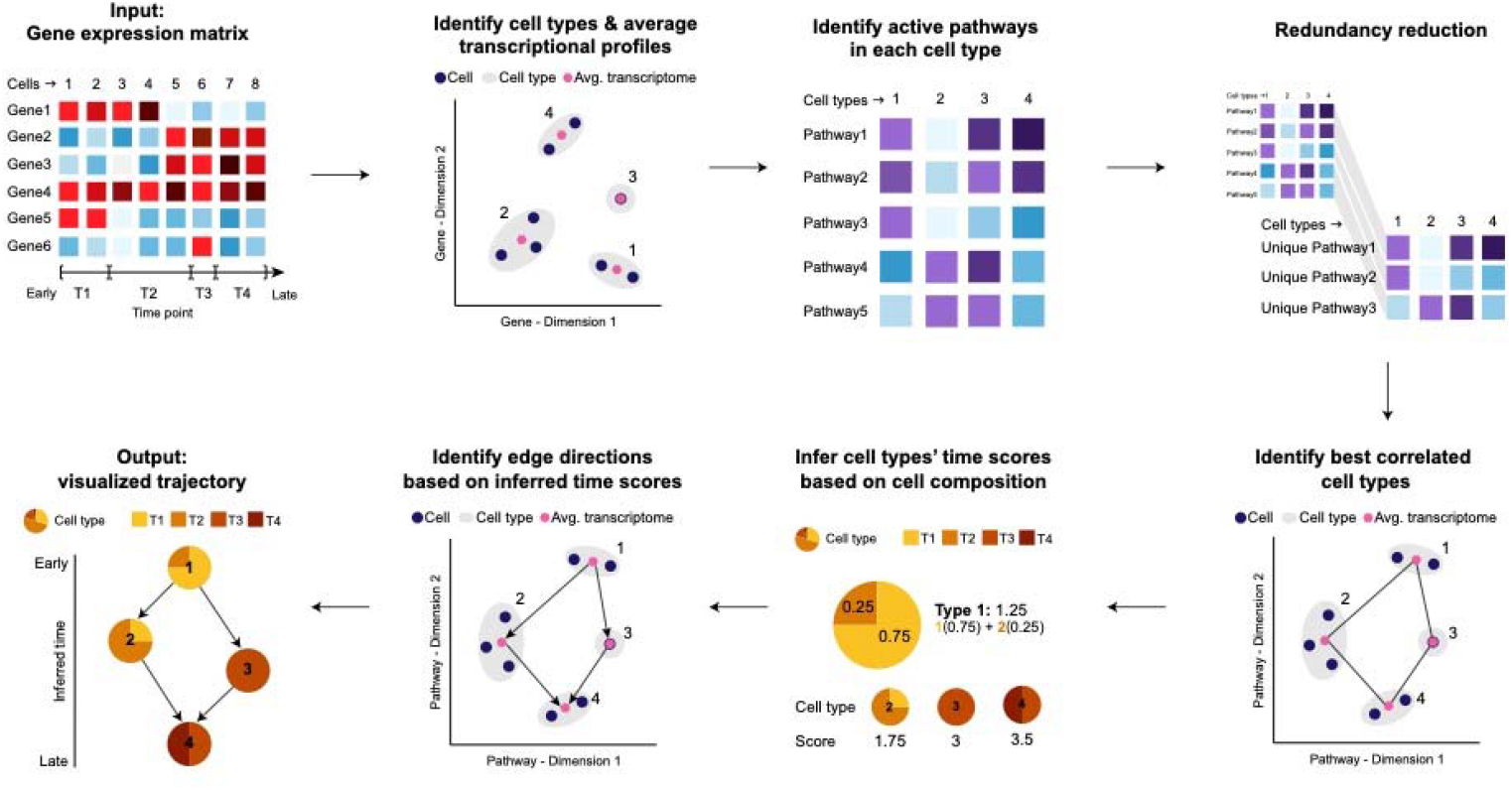
Schematic of the Tempora algorithm.

We abstract the trajectory as a network of cell clusters, where vertices represent the cell types or states identified as clusters and edges represent transitions between types or states. To infer this network, Tempora uses ARACNE^16^, an established algorithm that identifies cluster relationships using mutual information (MI) on the cluster pathway enrichment profiles. ARACNE filters the MI network using the data processing inequality to remove edges with the smallest MI in all triples, which helps remove indirect connections (Figure 1). After constructing the trajectory, Tempora uses available temporal information from the input data to determine edge directions. First, each cluster is assigned a temporal score corresponding to its cell composition from each time point, so that a cluster containing more cells from an early time point will have a low score and vice versa. Trajectory network edges are then directed so that their sources have a lower temporal score than their targets, indicating a transition from an early cell state to a later cell state. The trajectory is visualized using Sugiyama hierarchical layout algorithm^17^.

Tempora includes a downstream pathway exploration tool to determine and visualize pathways that change significantly over the trajectory. These pathways are identified by fitting a generalized additive model to the enrichment scores of each pathway across all clusters and selecting pathways whose expression patterns deviate significantly from the null model of uniform pathway enrichment scores across all time points.

### Validation on human skeletal muscle myoblast time-series data

We evaluated Tempora’s performance on human skeletal muscle myoblast (HSMM) data, which includes 271 cells collected at 0, 24, 48 and 72 hours after the switch of human myoblast culture from growth to differentiation media. The muscle myoblast culture is known to contain contaminating fibroblast cells, which originate from the same muscle biopsy used to establish the primary culture^9^. At the optimal clustering resolutions (see Methods), five clusters were identified and annotated with markers of proliferation (*CDK1*), muscle differentiation (*MYOG*) and contaminating myofibroblast cells (*SPHK1*)^9^ (Supplementary Figure 1a-d). Tempora identifies a branching trajectory connecting these clusters, rooted at the myoblast cluster that contains mostly cells at 0 hours after the media switch. This cluster leads to three separate branches, including a branch connected to the fibroblast cluster, one connected to the myotube cluster, and the last one connected to the partially differentiated myotube cluster via an intermediate cluster (Figure 2a). This branching trajectory agrees with the known biology of muscle differentiation *in vitro*, in which myoblasts proliferate and exit the cell cycle before differentiating into myotubes^9^. The fibroblast cluster contains equal proportions of cells from all time points and uniquely expresses myofibroblast markers (*SPHK1*). The equal numbers of cells from all time points in this cluster suggest that the contaminating cells were present in the earliest time point and persist in the culture over time, while its separation from the other two branches suggest that these cells do not participate in the differentiation process. Thus, Tempora identifies fibroblasts as a source of contamination in the myoblast culture, consistent with results from other trajectory inference methods^9,10^ and from the literature^18^. Another branch in this trajectory connects the myoblast cluster to the myotube cluster, which contains *MYOG*-positive cells mostly at 48 and 72 hours. (Figure 2a). *MYOG* is a required transcription factor for the terminal differentiation of myoblasts into myotubes and is rapidly upregulated when myoblasts start to differentiate around day 2 *in vitro*^19^. Therefore, the appearance of *MYOG*-positive myotubes at 48 hours and their connection to the myoblasts cluster, as predicted by Tempora, aligns with previous findings in the literature. Finally, the myoblast cluster is also connected to an intermediate cluster, which contains 75% cells from two early time points, expresses lower level of *CDK1* and does not express *MYOG* (Figure 2a). The low *CDK1* expression suggests that cells in this cluster have begun to exit the cell cycle to start differentiation, thus representing an intermediate state between proliferating myoblasts and differentiated muscles that is consistent with our understanding of muscle differentiation^20^. This intermediate cluster is predicted to give rise to a cluster of partially differentiated cells, which contains mostly cells from later time points and expresses low levels of the muscle-specific transcription factor *MYOG*. Since HSMM cultures have been noted to differentiate asynchronously and with less than 100% efficiency, cells in this partially differentiated cluster likely represent a cell population that is slower to differentiate or failed to go through differentiation as observed in previous studies^19^. Tempora, thus, predicts a branching trajectory that matches the structure and gene expression patterns of the known trajectory^20^.

**Figure 2.**
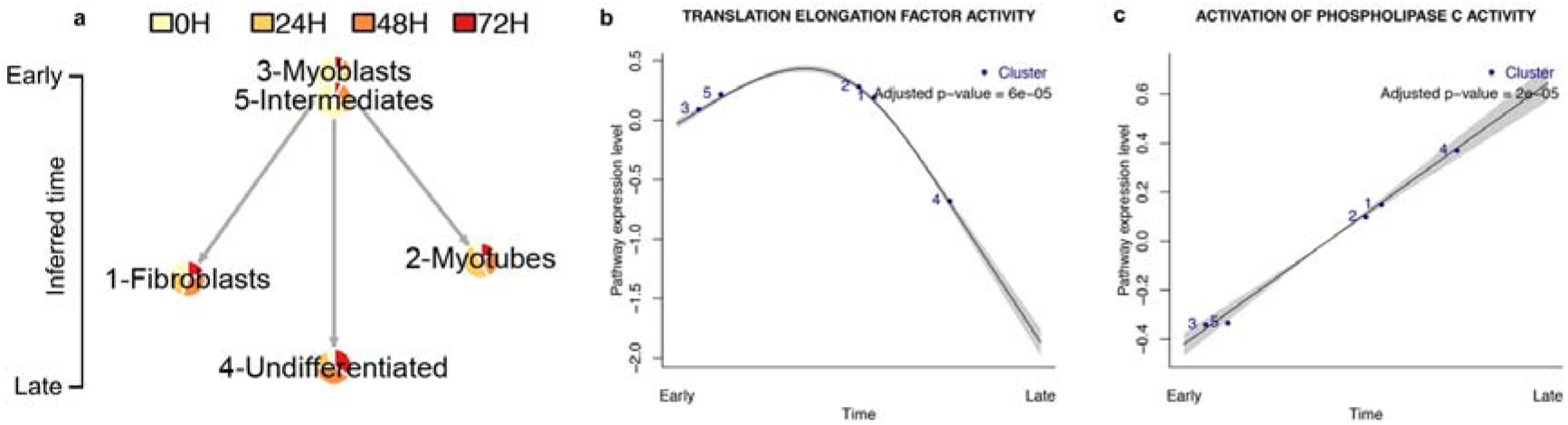
Tempora analysis of the HSMM dataset. **a.** Tempora trajectory built on clusters in the HSMM data set. **b-c.** Time-dependent pathways in the HSMM data set as discovered by the Tempora pathway exploration feature. Grey regions depict 95% confidence intervals.

We used the pathway exploration feature of Tempora to identify pathways whose enrichment changed over time. Pathways enriched early in the differentiation process include the cell cycle, biosynthesis and protein translation (Figure 2b-c). Pathways upregulated later are associated with the formation of myotubes, which include morphogenesis and phospholipase C signaling, which regulate myogenic activity^20,21^ (Figure 2b-d). Thus, Tempora’s pathway exploration component can be used downstream of trajectory inference to identify pathways with interesting activity profiles over the time-series.

### Validation on mouse cerebral cortex time-series data

We next applied Tempora on an embryonic murine cerebral cortex development scRNAseq data set, which contains approximately 6,000 neural cells collected at embryonic days 11.5 (E11.5), E13.5, E15.5 and E17.5^5^ (Figure 4a). These cells cover a wide spectrum of neuronal development, from the early precursors (apical precursors (APs) and radial precursors (RPs)) to intermediate progenitors (IPs) and differentiated cortical neurons. Data at all time points were aggregated and batch effects were corrected with Harmony before clustering (see Methods). We annotated the seven resulting clusters using marker genes for APs (*Sox2*, *Pax6*, *Hes1*, *Mki67*), RPs (*Edrnb*, *Vim*, *Slc1a3*), IPs (*Eomes, Gadd45g, Mfap4, Sstr2*), newborn neurons (*Tbr1, Tubb3, Foxp2, Reln*) and neurons (*Tubb3, Bhlhe22, Satb2, Fezf2, Mef2c, Gria2*) (Figure 4b-f). This resulted in the annotation of two AP/RP clusters mostly comprising cells at E11.5, which is consistent with the known emergence of RPs from APs at E11^5,22^, as well as two IP clusters, one IP/young neuron cluster and two neuron clusters, all of which contain cells from multiple timepoints as expected from their gradual specification over time^5^ (Supplementary Figure 2b-g).

Tempora predicts three trajectories, two rooted at the two AP/RP clusters and one rooted at an early IP cluster (Figure 3a). Each of the two AP/RP lineages has two branches: one terminating at an IP/young neuron cluster and another converging at a late neuron cluster. The lineage predicted by Tempora aligns with our understanding of AP/RP asymmetric division to generate IPs and neurons in early corticogenesis^5,22,23^. To better understand why there are two trajectories arising from two AP/RP clusters instead of one AP/RP cluster transforming into another AP/RP cluster in a single trajectory, we compared the gene expression profiles of the two clusters and identified cell cycle markers, such as *Mki67* and *Cdk1*, to be differentially expressed in one cluster versus the other. This suggests that the two AP/RP clusters differ based on their cell cycle state: one is actively proliferating and expressing cell cycle markers while the other is not (Supplementary Figure 2d), consistent with the known decreased proliferation of APs as they transition to RPs^5,24^. The observation that both AP/RP clusters contain equal proportion of cells from all time points suggest that these two proliferative and non-proliferative AP/RP populations arise before the time-series started, instead of one transforming into the other. Similarly, the IP cluster that serves as the root of the third trajectory contains many cells from the earliest time point and is thus unlikely to come from either of the AP/RP clusters, but may instead arise from earlier APs that are not captured in this time-series data. This IP cluster is predicted to give rise to a cluster of young neurons, which then mature into neurons as denoted by a dashed line in Figure 3a due to the high similarity in temporal scores between the young neurons and neurons cluster. These transitions predicted by Tempora are consistent with our understanding of neurogenesis^5,22^. Tempora, thus, accurately identifies distinct trajectories originating from different populations in the murine cerebral cortex development data.

**Figure 3.**
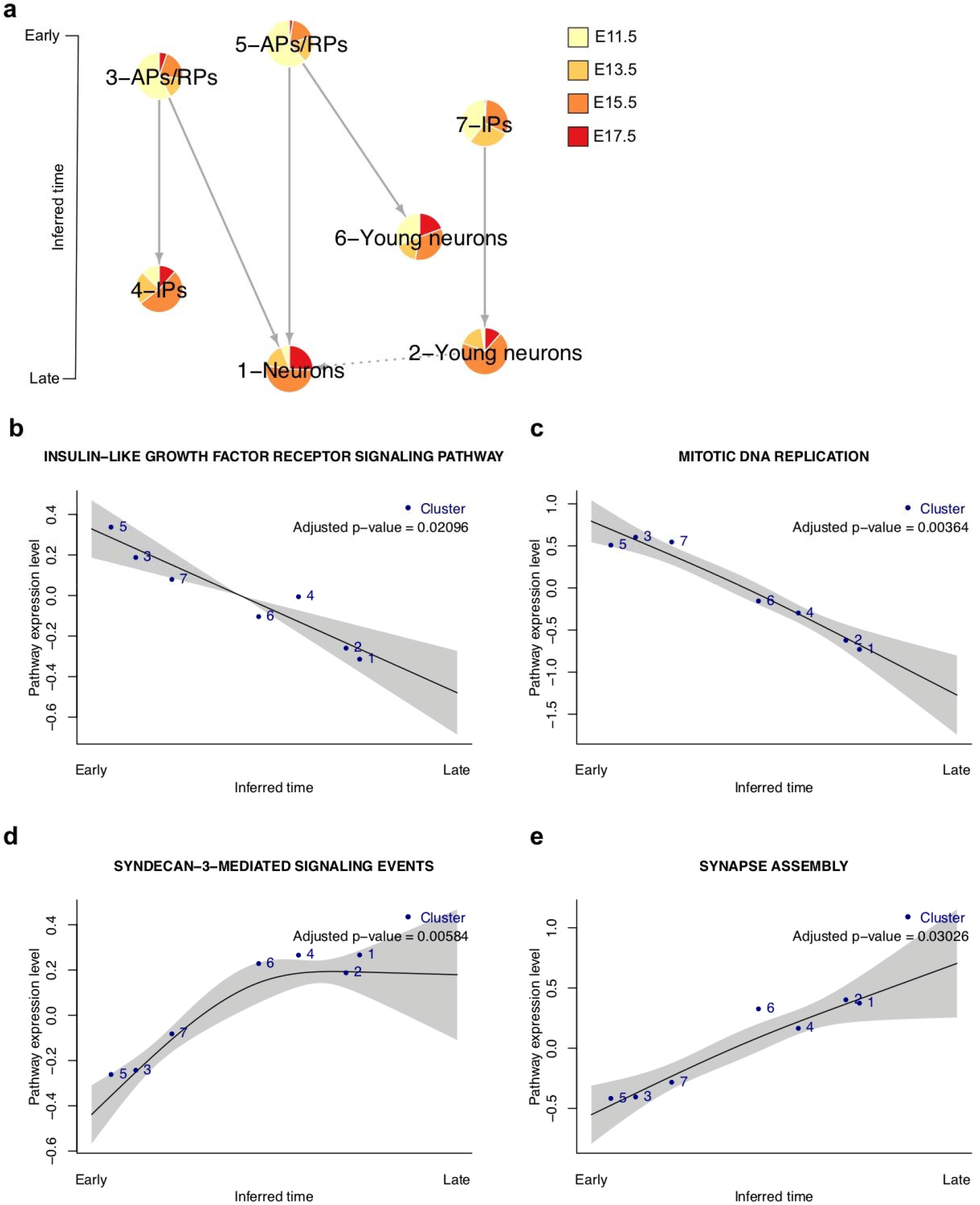
Tempora analysis of the murine cortex dataset. **a.** Tempora trajectory built on clusters in the murine cerebral cortex data set. **b-e.** Time-dependent pathways in the murine cerebral cortex data set as Tempora pathway exploration feature. Grey areas depict 95% confidence intervals.

We used the pathway exploration feature of Tempora to analyze time-dependent pathway activity levels in the data. DNA replication and other mitotic pathways are enriched early (Figure 3b), while neuron-related pathways, such as synapse activity, dendritic morphogenesis and neurotransmitter synthesis, are enriched later (Figure 3d-e). These patterns are consistent with the known proliferation of neural progenitors at the beginning of neurogenesis^24^ and neuronal activities of newborn neurons later in the process^25^. Tempora also identifies more subtle changes in signaling pathways over time, such as the early enrichment of Insulin-like growth factor signaling and the later upregulation of Syndecan-3 mediated signaling, both of which are consistent with their known roles in early neurogenesis and neural circuit assembly, respectively^26–28^ (Figure 3b-e).

### Performance evaluation

To evaluate Tempora’s performance and compare it to other cell trajectory inference methods, we measured the ability of a selected set of methods to recapitulate a gold standard set of known cell trajectories. For ease of comparison, we formalized all trajectories, both predicted and known, as graphs (networks), with nodes representing cell types, and directed edges representing parent-child relationships between connected nodes. We used two performance scores: graph edit distance (GED) or ‘mismatch score’, which measures the number of edge and node additions or removals required to transform the inferred trajectory to the known trajectory, and F1 score or ‘accuracy score’, which is the harmonic mean of precision and recall of gold standard directed edge identification.

We first constructed model trajectories for the *in vitro* differentiation of human myoblasts and murine cortical development through literature search^20,22,23,29^ (Figure 4a) and compare Tempora’s inferred trajectories to this gold standard. Human myoblasts, after exiting the cell cycle, transition through intermediate states before differentiating into myotubes^20,30^. Since myoblasts have varied differentiating potentials and rates, a portion of them will become myotubes while the rest remain undifferentiated, i.e. they do not, or have yet to, express myogenic transcription factors such as MYOG, which leads to two possible branches from the intermediate state(s)^19^. The starting culture, however, is often contaminated with fibroblast cells, which exert paracrine influence on the differentiation process but cannot differentiate into myotubes^31,32^. These contaminating cells, thus, form a branch separate from the main differentiation trajectory (Figure 4d).

**Figure 4.**
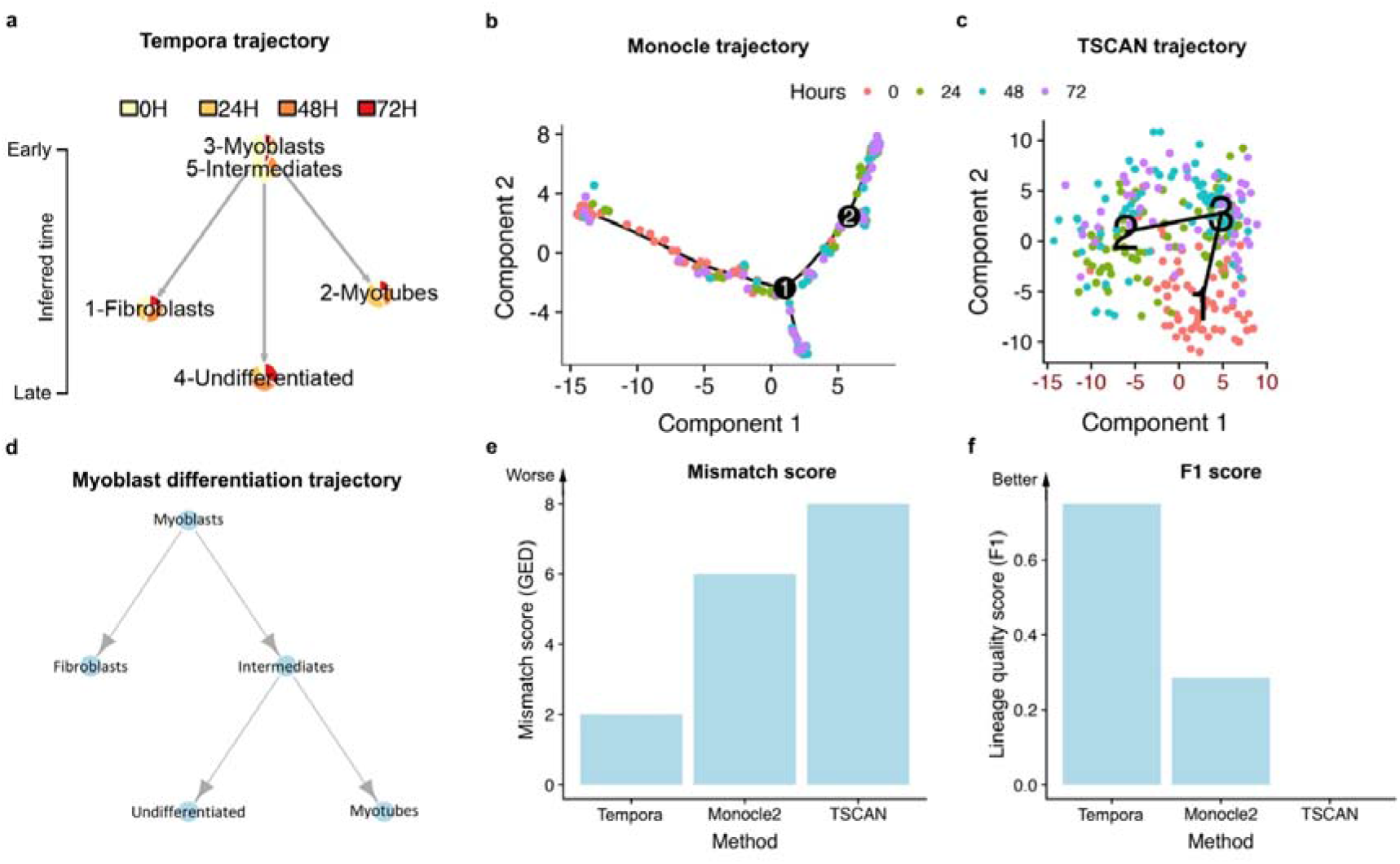
Performance evaluation on the HSMM data set. **a-c.** Trajectories of the HSMM data set inferred by **a.** Tempora, **b.** Monocle 2 and **c.** TSCAN. **d.** The model trajectory used to evaluate the accuracy of all inferred trajectories. **e.** Mismatch scores and **f.** accuracy scores of trajectories from the three evaluated methods.

Tempora’s predicted lineage (Figure 4a) is closely aligned with the model lineage, except it connects the myotubes cluster to the myoblasts instead of to the intermediate state. This results in a mismatch score of 2, which means that only two edges need to be changed in Tempora’s output to match the gold standard (Figure 4e). Furthermore, Tempora achieves a high accuracy score of 0.78 as it is able to infer the correct directions of most edges in the trajectory, except for the missing intermediate state to myotubes connection (Figure 4f). This result demonstrates that Tempora is able to infer a trajectory in the HSMM data set that is mostly consistent with the gold standard.

Murine corticogenesis consists of transitions between well-characterized cell types. The apical precursors (APs), which delaminate from the neuroepithelium, divide asymmetrically to self-renew and give rise to neurons^22^. At around E11, APs transition to radial precursors (RPs), which continue the asymmetric division to generate neurons either directly or indirectly through IPs^22,23^ (Figure 5d). Tempora’s inferred trajectory of the murine cerebral cortex data set achieves a low mismatch score and high accuracy score. It predicts almost all possible transitions between different cell types in the systems, only missing the IPs to neuron connection, which results in a mismatch score of 1 (Figure 5e). Despite this error, Tempora achieves a high accuracy score of 0.9 on the murine cerebral cortex data set, demonstrating that it can accurately identify directed connections between cell types in a large data set with multiple branches (Figure 5f).

**Figure 5.**
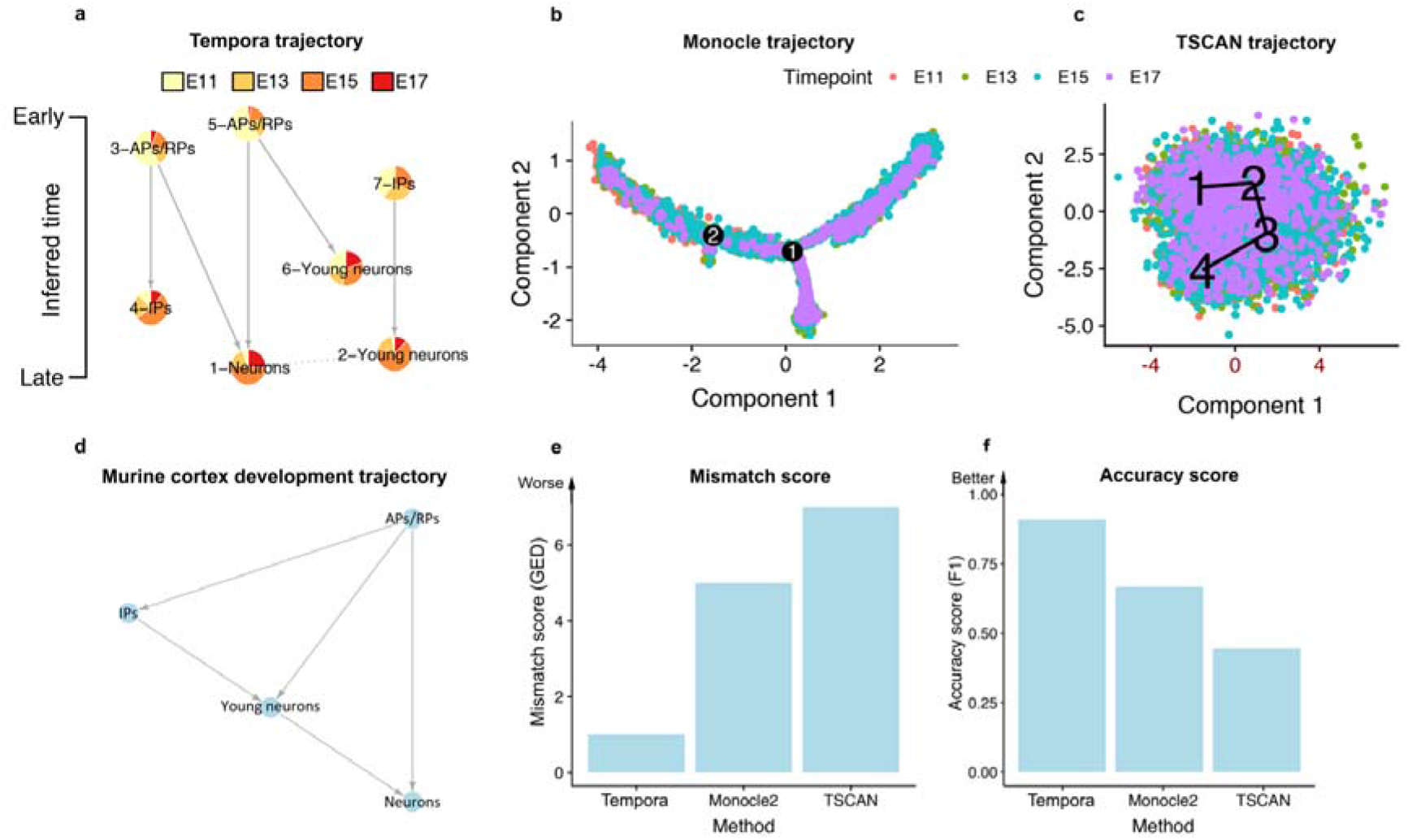
Performance evaluation on the murine cerebral cortex data set. **a-c.** Trajectories of the murine cerebral cortex data set inferred by **a.** Tempora, **b.** Monocle 2 and **c.** TSCAN. **d.** The model trajectory used to evaluate the accuracy of all inferred trajectories. **e.** Mismatch scores and **f.** accuracy scores of trajectories from the three evaluated methods.

We next compared Tempora’s performance with Monocle and TSCAN on the HSMM and murine cerebral cortex data sets when using each method’s default recommended protocols (see Methods). When applied to the HSMM data, Tempora outperforms both Monocle 2 and TSCAN (Figure 4). Monocle 2 predicts a trajectory with four states, annotated as myoblasts, partially differentiated, mesenchymal and differentiated myotubes, respectively. The trajectory branches at the end of the myoblast state into the other three states without a clear intermediate state (Figure 4b). This lack of intermediate state leads to Monocle’s worse mismatch score of 6. TSCAN’s trajectory has a linear structure, with the myoblasts at the root (state 1, annotated on the TSCAN plot) progressing through an intermediate state (state 3) and terminating at a cluster with mixed differentiated and undifferentiated cells (state 2). TSCAN’s trajectory is penalized because it neither separates the mesenchymal cells from the undifferentiated cells nor undifferentiated cells from differentiated myotubes at the terminal states, increasing its mismatch score to 8 (Figure 4c). Overall, even though all methods predict similar branching trajectories, Tempora performs best in terms of mismatch score as it correctly identifies the expected cell states and their directions in the HSMM data (Figure 4e). To calculate the F1 score on undirected trajectories inferred by Monocle 2 and TSCAN, we first determined the origin of the trajectories based on high expression of *CDK1*, *CCND5* for myoblasts in the HSMM data set^9^, then added directions to the inferred trajectories by directing all edges outward from the origin. Tempora’s accuracy of 0.78 outperformed Monocle 2 (accuracy of 0.3) and TSCAN (accuracy of 0) (Figure 4f).

Tempora also outperforms Monocle 2 and TSCAN on the murine cerebral cortex data set, which contains more transitions than the HSMM data set. Monocle 2 infers a branched lineage at two main branches: one from an APs/RPs branch to two IP branches (branchpoint 2) and one from the larger IP branch to two neuron branches (branchpoint 1) (Figure 5b). The early neurons are merged in both of the neuron branches instead of identified as a distinct state. Monocle 2’s trajectory is thus penalized for not identifying this young neuron state as well as its inability to predict the direct differentiation from APs/RPs to neurons, thus achieving a mismatch score of 5. TSCAN predicts a linear trajectory that connects APs/RPs to IPs, then to two neuron clusters (Figure 5c). TSCAN is penalized because it forces an erroneous connection between two neuron clusters, and similar to Monocle 2, it does not recognize a separate young neuron state and the direct differentiation link between APs/RPs to neurons. This results in a higher mismatch score of 7. Tempora’s mismatch score of 1, thus surpasses that of Monocle 2 and TSCAN by at least five times (Figure 5e). To calculate accuracy scores on trajectories from these methods, we used *Sox2* neural stem cell marker expression to infer that both Monocle 2 and TSCAN lineages are rooted at the branch with the highest number of E11.5 cells (leftmost branch in Monocle 2 trajectory and branch 1 of TSCAN trajectory) and determined that all edges go outward from this root. The inferred directions of both predicted lineages are consistent with the gold standard trajectory. Tempora (accuracy score of 0.9) significantly outperforms Monocle 2 (accuracy score of 0.66) and TSCAN (accuracy score of 0.44) on the murine cerebral cortex data set (Figure 5f).

### Comparison of Tempora with and without pathway enrichment analysis

To understand the impact of using pathway enrichment profiles on trajectory inference compared to the gene expression profile input used by other methods, we compared trajectories in the HSMM and murine cerebral cortex data sets using Tempora with and without the pathway enrichment analysis (PEA) step. Removing the PEA step resulted in poorer performance, as evident in an up to 4-fold increase in mismatch scores and a 3-fold decrease in accuracy scores (Figure 6).

**Figure 6.**
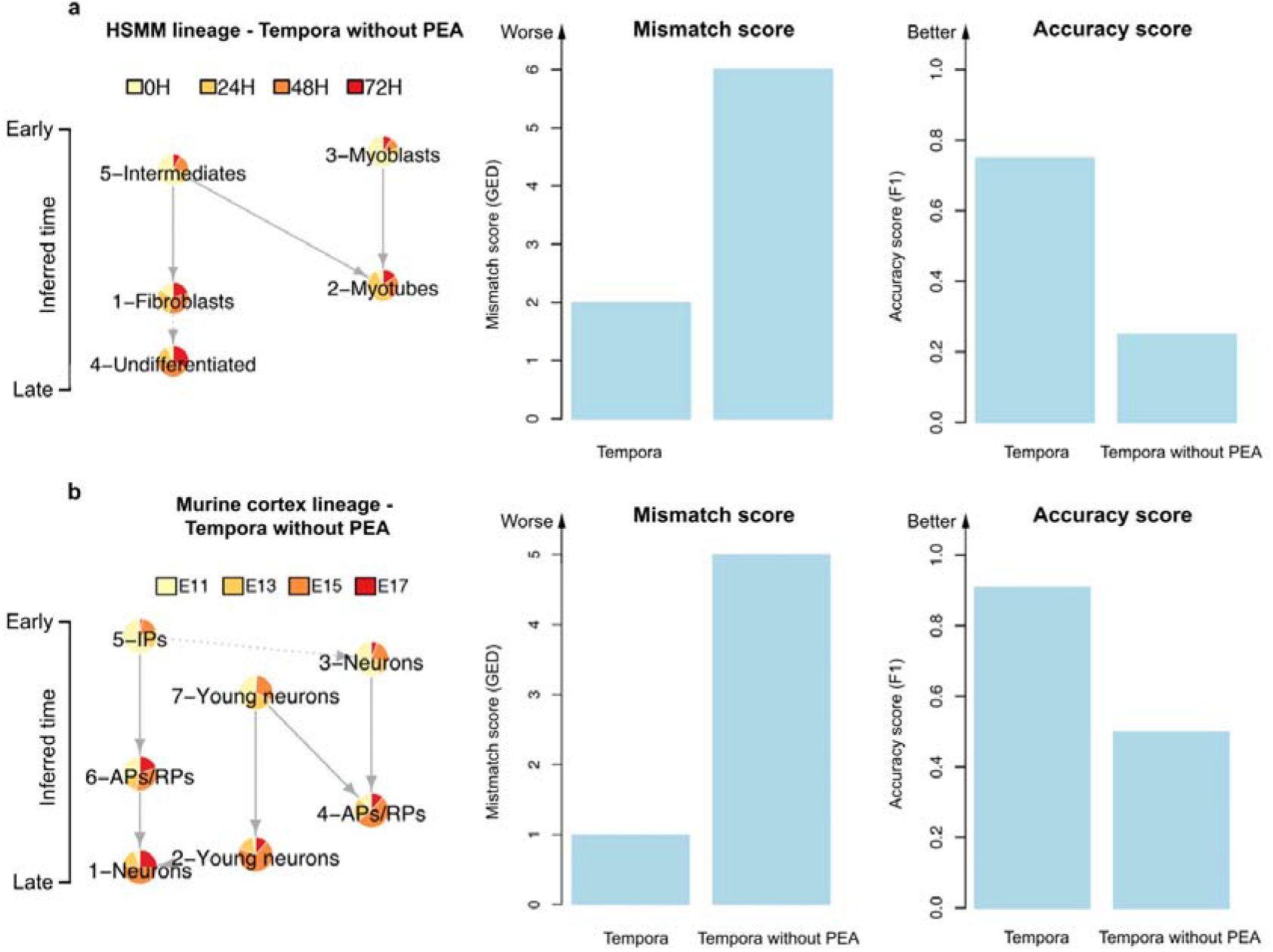
Performance evaluation of Tempora with and without pathway enrichment analysis (PEA). **a.** Performance of Tempora on HSMM and **b.** murine cerebral cortex data set with and without PEA.

Upon closer examination of the resulting trajectories, we observed that gene-input trajectories contain more edges between clusters with similar temporal scores compared to pathway-input trajectories, whose edges often connect clusters from different time points. We propose that this trend can be explained by the high similarity in gene expression profiles of clusters that are closer in developmental time, a fundamental assumption made by trajectory inference methods that rely on gene expression profile-based distance metrics to order cells. To test this hypothesis and better understand the discrepancies in inter-cluster gene vs. pathway enrichment profile similarity, we calculated the Pearson correlation between the gene and pathway enrichment profiles of all pairs of clusters in each data set. We found striking differences in the dynamic range of correlation observed: while correlations between gene expression profiles are uniformly strong and positive across all pairs of clusters, correlations between pathway enrichment profiles are negative for clusters of different cell types (neurons vs. APs, myoblasts vs. fibroblasts) and positive for clusters of the same cell types (neurons vs. neurons) (Supplementary Figure 3a-b). These differences suggest that the highly similar gene expression profiles across clusters make them less informative than pathway enrichment profiles in capturing changes along a trajectory.

## DISCUSSION

We have described and evaluated Tempora, a novel pathway-based cell trajectory inference method for time-series scRNAseq data. Tempora uses an information theoretic approach to build a trajectory at the cluster level based on the clusters’ pathway enrichment profiles, effectively connecting related cell types and states across multiple time points. Taking advantage of the available time information, Tempora infers the directions of all connections in a trajectory that go from early to late clusters. Evaluation on two time series scRNAseq data sets with known developmental trajectories (*in vitro* differentiation of human skeletal muscle myoblasts and *in vivo* early development of murine cerebral cortex) demonstrate that Tempora can accurately predict the lineages in time series data containing cell populations spanning all developmental stages, outperforming leading trajectory inference methods. Furthermore, downstream analyses using Tempora’s pathway exploration feature identifies pathways known to be important during the process under study, demonstrating the method’s ability to recapitulate and discover relevant biological signals during development processes.

Our analysis follows established scRNAseq analysis workflows, in which certain decisions can affect Tempora’s output. Tempora assumes that user input includes an optimized clustering solution for their data. If the clustering is not optimal, the output trajectory may be too general or too detailed. Over-clustering, when clusters are split too much (e.g. splitting a single cell type into two clusters), can lead to parallel edges originating from oversplit clusters and terminating at another cluster, ostensibly suggesting a multiple-parent lineage (Supplementary Figure 4). Under-clustering can result in overly simplified lineages (Supplementary Figure 4). Under-clustering can also lead to certain cell types appearing at one time point but absent from another, because they have been clustered with other cell types at one time point and not the other. These challenges are inherent to clustering high-dimensional scRNAseq data, which relies on user-input parameters to determine the number of clusters, and no gold standard exists to guide the selection of these parameters^33^.

An important part of Tempora’s analysis is batch effect assessment and correction, as time-series data are often collected in batches, thus easily subjected to technical variations between experimental runs. We used Harmony to correct for batch effect in the two scRNAseq datasets used in this study before downstream trajectory analysis. Without such correction, Tempora’s performance decreased slightly on both gold standards (Supplementary Figure 5, Supplementary Figure 6). This is likely due to the suboptimal clustering driven by batch effects, which results in less accurate inference of trajectories based on the resulting clusters. Thus, we recommend running batch correction and data alignment procedure before Tempora analysis.

The increasing use of time-series scRNAseq to investigate dynamic biological processes, including development and differentiation, present both opportunities and challenges. The larger cell numbers and types captured in time-course experiments enable researchers to discover rare cell types and study cell transitions with higher resolution, yet the non-synchronous nature of the populations across time points present a computational challenge to automatically infer cell trajectories^34^. Using time information to supervise the trajectory inference process enables accurate identification of cell types consisting of cells from different time points as well as the lineage connections between them. When combined with other methods to infer population and transcriptional dynamics to analyze time-series scRNAseq data, time-series based analysis can generate powerful insights into dynamic processes and their biological regulation.

## METHODS

### Single-cell RNAseq data

Two time-course scRNAseq data sets were used to validate Tempora, one on the *in vitro* differentiation of human skeletal muscle myoblasts (HSMM) and the other on an *in vivo* sample of early murine cerebral cortex development. HSMM read count data was accessed from GEO, accession number GSE52529^9^. Murine cerebral cortex data were downloaded from GEO, accession number GSE107122^5^.

Both data sets were filtered to remove lowly expressed genes (defined as those found in less than 3 cells) and damaged cells with high mitochondrial genome transcript content (4 median absolute deviations above the median). After this initial filtering step, the murine cerebral cortex data were further filtered to remove non-cortical cells, as done in the original publication^5^. These included cells expressing *Aif1* (microglia), hemoglobin genes (blood cells), collagen genes (mesenchymal cells), as well as *Dlx* transcription factors and/or interneuron genes (ganglionic eminence-derived cells)^5^. The data sets were then normalized using the deconvolution method implemented in the *scran* R package, which pools cells with similar gene expression profiles and library sizes together to normalize^35^. Afterwards, cells were iteratively clustered in Seurat at increasing resolutions until the number of differentially expressed genes between two neighboring clusters reached 0, as determined by scClustViz^36^. We then chose the optimal clustering resolution, defined as the point where the number of clusters was maximized while the number of DE genes between neighboring clusters remained larger than 0, and annotated all clusters by examining expression of known marker genes using scClustViz^36^. The resulting clusters represent cell types or states that are stable over the developmental process, such as apical progenitor cells in murine cerebral cortex development and myoblasts in muscle development.

### Data preprocessing, batch effect correction and clustering

Tempora takes processed scRNAseq data as input, either as a gene expression matrix with separate time and cluster labels for all cells, or a Seurat object containing gene expression data and a clustering result. Tempora does not implement clustering or batch effect correction as part of its pipeline and assumes that the user has input a well-annotated cluster solution free of batch effect into the method.

Since a good clustering result is central to the successful application of Tempora on a data set, we recommend users take advantage of methods such as scClustViz^36^ to visualize clusters at different resolutions, analyze cluster relationships across resolutions as well as investigate marker gene expression to help choose appropriate clustering parameters. Furthermore, users should define a set of rules that they use to determine the optimal clustering resolution to maintain consistency between analyses^36^.

### Pathway enrichment analysis

Tempora calculates the average gene expression over all cells in a cluster for all clusters as input by the users and determines the pathway enrichment profile of each cluster using Gene Set Variation Analysis (GSVA)^15^. By analyzing scRNAseq data on the cluster level instead of the single-cell level, Tempora amplifies gene expression signals from similar cells in a cluster to alleviate the typical problem of low sensitivity per cell of popular scRNA-seq experimental methods, as well as to reduce the number of nodes in the inferred lineage, allowing users to interpret it more easily. The default pathway gene set database Tempora uses is the Bader Lab pathway gene set database without electronic annotation Gene Ontology terms (Human_GO_AllPathways_no_GO_iea_August_01_2019_symbol.gmt, accessed at http://download.baderlab.org/EM_Genesets/current_release/Human/symbol/Human_GO_AllPathways_no_GO_iea_August_01_2019_symbol.gmt, and Mouse_GO_AllPathways_no_GO_iea_August_01_2019_symbol.gmt, accessed at http://download.baderlab.org/EM_Genesets/current_release/Mouse/symbol/Mouse_GO_AllPathways_no_GO_iea_August_01_2019_symbol.gmt), filtered to include gene sets between 10 and 500 genes in size^37^. The enrichment scores of all P pathways in each cluster make up the cluster’s pathway enrichment profile, which is a vector of length P.

Since pathway gene set databases contain redundant pathways and this redundancy is not evenly distributed across the database (e.g. well studied pathways are better represented), Tempora uses PCA to reduce redundancy in all pathway enrichment profiles and identifies the top *n* principal components, defined as the components before the slope levels off (“the elbow”), in a scree plot, to input to downstream trajectory construction steps. In this study, 5 PCs were used for analysis of the HSMM data and 6 for the murine cortex data.

### Filtered mutual information network

We conceptualize the cell (cluster) trajectory from different timepoints of a developmental process as a graph (network), where vertices represent clusters and edges represent parent-child relationships between these vertices. Tempora employs the mutual information (MI) rank approach implemented in ARACNE^16^ to calculate MI between all cluster pairs present in the data. The data-processing inequality is then applied by ARACNE to remove the edge with the lowest MI in each triple to reduce the number of indirect interactions between clusters. This results an undirected network where nodes are cell clusters and weighted edges represent MI strength relationships between clusters.

### Direction identification

Tempora uses time information to determine the edge directions in the constructed MI network. Tempora assigns each time point a sequential, ordinal value corresponding with its distance from the earliest time point and calculates a temporal score for each cluster based on its composition of cells from each timepoint. Specifically, the temporal score, *T*_*k*_, of cluster *k* consisting of *p*_*i*_ percent cells at timepoint *i* is calculated as:

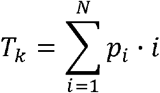

Where *N* is the number of time points. Under the assumption that cell differentiation progresses unidirectionally from stem or progenitor cells (early timepoints) to differentiated (late timepoints) cells, Tempora assigns directions to all edges in the network so that edges point from clusters with low temporal scores to clusters with high temporal scores. For edges that connect clusters with similar temporal scores (with the similarity threshold defined by the users), Tempora does not assign directions as these edges can represent small transitions in cell states over a short time, in which the unidirectional assumption may not hold.

### Identification of time-dependent pathways

Tempora identifies pathways that vary over time by fitting a generalized additive model on the pathway enrichment scores of each pathway across all clusters over time and using ANOVA to compare the fitted model with the null model of uniform pathway enrichment over time. Pathways with adjusted p-values below a user-defined threshold, with a default value of 0.05, are reported as significantly varying over time. The model fitting and statistical testing are done using the *mgcv* package in R.

### Evaluation and comparison with other methods

We evaluated Tempora by comparing its predicted cell lineages to known lineages manually curated from the literature. The lineages are represented as graphs, with vertices representing cell types and edges representing lineage connections. Two approaches were then used to measure the accuracy of Tempora and other methods in predicting the known lineage: a mismatch score, implemented as the graph edit distance (GED), and accuracy, implemented as the F1 score.

### Model trajectory construction

We manually curated the model trajectories for the *in vitro* differentiation of human myoblasts and murine cortical development through literature search^20,22,23,29^ and represented the lineage relationships between different cell types in the system as a graph. Each node in a model trajectory represents a distinct cell type as noted in the literature and described in the Cell Ontology^38^, while the edges represent lineage connections (*develops_from* relationship in the Cell Ontology) between these cell types.

### Mismatch score

We used the unweighted GED metric to measure the number of mismatches between the predicted and known trajectory, both formalized as undirected graphs to enable comparisons with methods that do not predict edge directions^39^. GED is formally defined as the smallest total number of graph edit operations needed to transform one graph into another. In this context, the permitted operations included insertion and deletion of edges or vertices.

To calculate the mismatch score between a pair of graphs, we first label each cluster in the inferred trajectory with the cell type(s) it contains, based on expression of a set of well-known marker genes. The cell types used for labeling are standard terms from the Cell Ontology database. If multiple clusters contain one cell type, they are assigned the same label. In case one cluster contains multiple cell types as defined by the positive expression of multiple marker genes, we label the cluster with both cell types. We then calculated the number of differences in the cell types of the predicted and known trajectories, as well as in the adjacency matrices of both trajectories. The sum of these two differences is the mismatch score for each pair of graphs.

### Accuracy score

To compare the accuracy of Tempora’s time-based direction inference with the model trajectory, we calculated the F1 score on each predicted trajectory as follows:

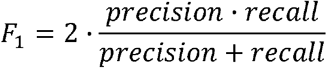

in which true positives (TP) are edges present in both the model and the predicted trajectory, false positives (FP) are edges in the predicted trajectory but not in the model, and false negatives (FN) are edges in the model but not in the predicted trajectory. An edge in the predicted graph is considered true positive only when its two vertices and direction match those of an edge in the model graph.

To calculate the accuracy score on undirected trajectories inferred by Monocle 2 and TSCAN, we first determined the origin of the trajectories based on high expression of a set of marker genes (*CDK1*, *CCND5* for myoblasts in the HSMM data set^9^ and *Sox2* for apical precursors in the mouse cerebral cortex data set^5^), then added directions to the inferred trajectories by directing all edges to go outward from the origin.

### Monocle

We applied Monocle 2^40^ on two data sets in this study using the method’s recommended protocol. Genes used for the pseudotime inference process were determined using the dpFeature procedure, in which cells were first clustered and the 1000 most differentially expressed genes across the clusters were selected for downstream analyses. To formalize a Monocle trajectory as a graph, we considered each state, or segment of the tree, as a vertex, and connected the vertices with appropriate edges to recapitulate Monocle’s output.

### TSCAN

We applied TSCAN^10^ on two data sets in this study using the Shiny GUI, which allowed us to make use of additional marker gene visualization features not available with the TSCAN command line tool. As TSCAN clusters each data set and constructs a minimum spanning tree (MST) on the clusters, its output lends itself well to the graph formalization we use. We optimized the number of clusters used in trajectory construction using TSCAN’s built in optimization feature. We then retained the cluster-level MST that TSCAN outputs for each data set and considered each cluster a vertex, while the segments of the MST are edges in the graphs we use. We then determined the roots and directions of the graph as described in the section on accuracy score calculation.

## DECLARATIONS

### Availability

The Tempora R package and all validation data used in this study can be accessed at https://github.com/BaderLab/Tempora.

### Authors’ contributions

T.N.T and G.D.B conceptualized the work, interpreted the results, drafted and revised the manuscript. T.N.T processed the single-cell scRNAseq data, developed the algorithm and prepared the R package. G.D.B supervised the study.

## Acknowledgements

We thank M. Abou Chakra and B. Innes for helpful discussion and feedback on this manuscript.

## REFERENCES

1 Hwang, B., Lee, J. H. & Bang, D. Single-cell RNA sequencing technologies and bioinformatics pipelines. Experimental & Molecular Medicine 50, 96, doi:10.1038/s12276-018-0071-8 (2018).

2 Trapnell, C. Defining cell types and states with single-cell genomics. Genome Research 25, 1491–1498, doi:10.1101/gr.190595.115 (2015).

3 Kumar, P., Tan, Y. & Cahan, P. Understanding development and stem cells using single cell-based analyses of gene expression. Development 144, 17, doi:10.1242/dev.133058 (2017).

4 Weinreb, C., Wolock, S., Tusi, B. K., Socolovsky, M. & Klein, A. M. Fundamental limits on dynamic inference from single-cell snapshots. Proceedings of the National Academy of Sciences 115, E2467–E2476, doi:10.1073/pnas.1714723115 (2018).

5 Yuzwa, S. A. et al. Developmental emergence of adult neural stem cells as revealed by single-cell transcriptional profiling. Cell reports 21, 3970–3986 (2017).

6 Wang, P. et al. Dissecting the Global Dynamic Molecular Profiles of Human Fetal Kidney Development by Single-Cell RNA Sequencing. Cell Reports 24, 3554–3567.e3553, doi:https://doi.org/10.1016/j.celrep.2018.08.056 (2018).

7 Cui, Y. et al. Single-Cell Transcriptome Analysis Maps the Developmental Track of the Human Heart. Cell reports 26, 1934–1950.e1935 (2019).

8 Saelens, W., Cannoodt, R., Todorov, H. & Saeys, Y. A comparison of single-cell trajectory inference methods. Nature Biotechnology, doi:10.1038/s41587-019-0071-9 (2019).

9 Trapnell, C. et al. The dynamics and regulators of cell fate decisions are revealed by pseudotemporal ordering of single cells. Nature Biotechnology 32, 381, doi:10.1038/nbt.2859 (2014).

10 Ji, Z. & Ji, H. TSCAN: Pseudo-time reconstruction and evaluation in single-cell RNA-seq analysis. Nucleic Acids Research 44, e117–e117, doi:10.1093/nar/gkw430 (2016).

11 Street, K. et al. Slingshot: cell lineage and pseudotime inference for single-cell transcriptomics. BMC genomics 19, 477–477, doi:10.1186/s12864-018-4772-0 (2018).

12 Grün, D. et al. De Novo Prediction of Stem Cell Identity using Single-Cell Transcriptome Data. Cell Stem Cell 19, 266–277, doi:https://doi.org/10.1016/j.stem.2016.05.010 (2016).

13 Wolf, F. A. et al. PAGA: graph abstraction reconciles clustering with trajectory inference through a topology preserving map of single cells. Genome Biology 20, 59, doi:10.1186/s13059-019-1663-x (2019).

14 Bar-Joseph, Z., Gitter, A. & Simon, I. Studying and modelling dynamic biological processes using time-series gene expression data. Nature Reviews Genetics 13, 552, doi:10.1038/nrg3244 (2012).

15 Hänzelmann, S., Castelo, R. & Guinney, J. GSVA: gene set variation analysis for microarray and RNA-seq data. BMC bioinformatics 14, 7 (2013).

16 Margolin, A. A. et al. ARACNE: An Algorithm for the Reconstruction of Gene Regulatory Networks in a Mammalian Cellular Context. BMC Bioinformatics 7, S7, doi:10.1186/1471-2105-7-s1-s7 (2006).

17 Sugiyama, K., Tagawa, S. & Toda, M. Methods for Visual Understanding of Hierarchical System Structures. IEEE Transactions on Systems, Man, and Cybernetics 11, 109–125, doi:10.1109/TSMC.1981.4308636 (1981).

18 Hannon, K., Kudla, A. J., McAvoy, M. J., Clase, K. L. & Olwin, B. B. Differentially expressed fibroblast growth factors regulate skeletal muscle development through autocrine and paracrine mechanisms. The Journal of Cell Biology 132, 1151, doi:10.1083/jcb.132.6.1151 (1996).

19 Owens, J., Moreira, K. & Bain, G. Characterization of primary human skeletal muscle cells from multiple commercial sources. In Vitro Cell Dev Biol Anim 49, 695–705, doi:10.1007/s11626-013-9655-8 (2013).

20 Chal, J. & Pourquié, O. Making muscle: skeletal myogenesis in vivo and in vitro. Development 144, 2104–2122, doi:10.1242/dev.151035 (2017).

21 Faenza, I. et al. Expression of phospholipase C beta family isoenzymes in C2C12 myoblasts during terminal differentiation. Journal of Cellular Physiology 200, 291–296, doi:10.1002/jcp.20001 (2004).

22 Dwyer, N. D. et al. Neural Stem Cells to Cerebral Cortex: Emerging Mechanisms Regulating Progenitor Behavior and Productivity. The Journal of Neuroscience 36, 11394–11401, doi:10.1523/jneurosci.2359-16.2016 (2016).

23 Martynoga, B., Drechsel, D. & Guillemot, F. Molecular control of neurogenesis: a view from the mammalian cerebral cortex. Cold Spring Harbor perspectives in biology 4, a008359, doi:10.1101/cshperspect.a008359.

24 Homem, C. C. F., Repic, M. & Knoblich, J. A. Proliferation control in neural stem and progenitor cells. Nature reviews. Neuroscience 16, 647–659, doi:10.1038/nrn4021 (2015).

25 He, Z. & Yu, Q. Identification and characterization of functional modules reflecting transcriptome transition during human neuron maturation. BMC genomics 19, 262–262, doi:10.1186/s12864-018-4649-2 (2018).

26 Yu, C., Griffiths, L. R. & Haupt, L. M. Exploiting Heparan Sulfate Proteoglycans in Human Neurogenesis-Controlling Lineage Specification and Fate. Front Integr Neurosci 11, 28–28, doi:10.3389/fnint.2017.00028 (2017).

27 Nieto-Estévez, V., Defterali, Ç. & Vicario-Abejón, C. IGF-I: A Key Growth Factor that Regulates Neurogenesis and Synaptogenesis from Embryonic to Adult Stages of the Brain. Front Neurosci 10, 52–52, doi:10.3389/fnins.2016.00052 (2016).

28 Hsueh, Y.-P. & Sheng, M. Regulated Expression and Subcellular Localization of Syndecan Heparan Sulfate Proteoglycans and the Syndecan-Binding Protein CASK/LIN-2 during Rat Brain Development. The Journal of Neuroscience 19, 7415–7425, doi:10.1523/jneurosci.19-17-07415.1999 (1999).

29 van der Ven, P. F. M. et al. Differentiation of human skeletal muscle cells in culture: maturation as indicated by titin and desmin striation. Cell Tissue Res. 270, 189–198, doi:10.1007/BF00381893 (1992).

30 Bentzinger, C. F., Wang, Y. X. & Rudnicki, M. A. Building Muscle: Molecular Regulation of Myogenesis. Cold Spring Harbor Perspectives in Biology 4, doi:10.1101/cshperspect.a008342 (2012).

31 Hannon, K., Kudla, A. J., McAvoy, M. J., Clase, K. L. & Olwin, B. B. Differentially expressed fibroblast growth factors regulate skeletal muscle development through autocrine and paracrine mechanisms. The Journal of cell biology 132, 1151–1159 (1996).

32 Scata, K. A., Bernard, D. W., Fox, J. & Swain, J. L. FGF Receptor Availability Regulates Skeletal Myogenesis. Experimental Cell Research 250, 10–21, doi:https://doi.org/10.1006/excr.1999.4506 (1999).

33 Kiselev, V. Y., Andrews, T. S. & Hemberg, M. Challenges in unsupervised clustering of single-cell RNA-seq data. Nature Reviews Genetics 20, 273–282, doi:10.1038/s41576-018-0088-9 (2019).

34 Ding, J. et al. Reconstructing differentiation networks and their regulation from time series single-cell expression data. Genome research 28, 383–395, doi:10.1101/gr.225979.117.

35 L. Lun, A. T., Bach, K. & Marioni, J. C. Pooling across cells to normalize single-cell RNA sequencing data with many zero counts. Genome Biology 17, 75, doi:10.1186/s13059-016-0947-7 (2016).

36 Innes, B. & Bader, G. scClustViz - Single-cell RNAseq cluster assessment and visualization F1000Research 7, doi:10.12688/f1000research.16198.2 (2019).

37 Reimand, J. et al. Pathway enrichment analysis and visualization of omics data using g:Profiler, GSEA, Cytoscape and EnrichmentMap. Nature Protocols 14, 482–517, doi:10.1038/s41596-018-0103-9 (2019).

38 Bard, J., Rhee, S. Y. & Ashburner, M. An ontology for cell types. Genome biology 6, R21–R21, doi:10.1186/gb-2005-6-2-r21 (2005).

39 Gao, X., Xiao, B., Tao, D. & Li, X. A survey of graph edit distance. Pattern Analysis and Applications 13, 113–129, doi:10.1007/s10044-008-0141-y (2010).

40 Qiu, X. et al. Reversed graph embedding resolves complex single-cell trajectories. Nature Methods 14, 979–982, doi:10.1038/nmeth.4402 (2017).

